# So similar, yet so different: α-synuclein recognition of the isoform-specific VDAC protein–lipid interface

**DOI:** 10.64898/2026.10.13.758914

**Authors:** Tatiana K. Rostovtseva, Megha Rajendran, William M. Rosencrans, María Queralt-Martín, Mya S. Wolfe, Nina A. Bautista, Bethel G. Beyene, Motahareh G. Larimi, Alexander J. Sodt, Sergey M. Bezrukov

## Abstract

1.

In mammals, VDAC exists as three isoforms – VDAC1, VDAC2, and VDAC3 – each characterized by a distinct tissue-dependent distribution and physiologic role. Isoform-specific knockout studies in cells and mice demonstrate distinct physiological roles for each VDAC isoform. The isoforms share about 70% sequence homology and a conserved β-barrel structure and, when reconstituted into planar lipid membranes, form similar highly conductive, anion-selective, voltage-gated channels. A natural question, then, is how relatively minor differences in isoform sequences lead to striking differences in their interactions with protein partners and, consequently, in their physiological functions. To address this question, we used a small, intrinsically disordered cytosolic protein α-synuclein (αSyn) as a sensitive molecular probe. A comparative study with recombinant VDACs reconstituted into planar lipid membranes shows that, depending on the isoform, the αSyn-induced blockage exhibits up to 100-fold differences in on-rates and equilibrium constants. The isoforms also display distinct translocation regimes for αSyn, as evidenced by altered dependences of the off-rates on the applied voltage. Deletion of cysteine residues has a significant effect on αSyn binding. Combined with corresponding data obtained for VDAC2 substates – which show an order of magnitude variation in on-rates while maintaining the conserved equilibrium constants – and the results of molecular dynamics simulations, these findings point to a previously unrecognized step in VDAC inhibition by αSyn: an isoform-specific αSyn interaction with the VDAC protein-lipid interface. This finding opens a new paradigm for understanding VDAC isoform functions and developing isoform-specific therapeutics.

**New & Noteworthy:** Comparative analysis of α-synuclein interaction with three mammalian VDAC isoforms, complemented by corresponding data for VDAC2 substates and results from molecular dynamics simulations, reveals a new step in the chain of events leading to VDAC blockage. Specifically, our findings point to the isoform-specific interaction of αSyn with VDAC protein-lipid interface, thereby guiding a new strategy in the search for VDAC-targeting drugs.

## 2. Introduction

### VDAC is a mitochondrial communication hub

VDAC is recognized as a key pathway for most water-soluble mitochondrial metabolites across the mitochondrial outer membrane (MOM), including the primary energy currency ATP, other mitochondrial respiratory substrates, and small ions such as Ca^2+^. This is in contrast with more than 50 highly substrate-specific mitochondrial carriers in the mitochondrial inner membrane (IM), which also transport water-soluble substrates and ions across the IM to maintain mitochondrial function. VDAC is not only a unique conduit for metabolite and small-ion fluxes in and out of mitochondria, but it also dynamically regulates these fluxes (see, e.g., (1–6). Critically, VDAC actively regulates ATP flux through its characteristic voltage-gating mechanism (7).

The importance of this relatively simple channel, formed by 19 β-strands, arises from its position at the juncture between the mitochondria and the cytosol (3, 6), allowing VDAC simultaneous access to compartmentalized mitochondrial and cytosolic proteins and to the crosstalk between mitochondria and organelles such as the endoplasmic and sarcoplasmic reticulum (ER, SR) and lysosomes (8–10). VDAC has been found to interact with numerous proteins, such as different pro- and anti-apoptotic Bcl-2 family proteins (11–15), multiple kinases (16–19), cytoskeleton (20–22), neuronal (23–25), and mitochondrial proteins (26, 27). VDAC’s regulatory role over the interface between mitochondria and the cytosol makes it an important cellular communication hub. Therefore, it is not surprising that VDAC has been implicated in a wide range of mitochondria-associated pathologies, such as different types of cancer, diabetes, cardiovascular and neurodegenerative disorders such as Parkinson’s (PD) and Alzheimer’s disease (AD) (28–31). Changes in VDAC expression levels controlled either at the transcriptional level or through degradation of VDAC have been reported in many diseases (32, 33). In addition, the alteration of VDAC interaction with its interactome may also lead to mitochondrial dysfunction, for example, the increase in VDAC interaction with α-synuclein (αSyn) in PD (34) and with dimeric tubulin in cancer (35). Consequently, VDAC has emerged recently as a promising pharmacological target (36).

VDAC is a 33 kDa protein, which forms a monomeric, slightly anion-selective, relatively large channel (cut-off < 2 kDa) that allows passive diffusion of ions and small metabolites. There are three mammalian VDAC isoforms, each encoded by an individual gene: VDAC1, VDAC2, and VDAC3 (37, 38). VDAC isoforms show differences in tissue expression profiles, with VDAC1 and VDAC2 having the highest expression level (∼90%) in most of the tissues, especially in liver, brain, and heart (39), and VDAC3 the lowest (∼ 10%), except for testis (∼41%) (39–41). Each VDAC isoform has a unique physiological role, which was shown using VDAC isoform-specific embryonic knockout (KO). Namely, the knockout of VDAC1 (40) in mice causes only mild growth retardation and mitochondrial respiration deficiency (42), while VDAC2 KO is lethal or partially lethal in mouse embryos (12, 15) and VDAC3 KO in mice results in male infertility (43). VDAC2 is required for BAX/BAK-induced apoptosis (12, 15, 44), and VDAC1 plays a role in Ca^2+^-induced apoptosis in mitochondria (45), but VDAC3 has not yet been found to significantly impact apoptosis, suggesting isoform-specific roles in cell survival.

### VDAC isoforms are structurally and functionally similar

There is 60-70% sequence homology between the three VDAC isoforms (39). The crystal structure at the atomic level of resolution has so far been solved only for VDAC1 (46–48). The Cryo-EM structure of VDAC2 (27) and the predicted structure of VDAC3 show very similar 19-stranded β-barrels with a double α-helical N-terminus lying about in the middle of the pore (**Figure 1**). One difference between isoforms is in the number of cysteines: 2 in human VDAC1, 9 in VDAC2, and 6 in VDAC3, and the 11-residue N-terminal extension in VDAC2 (39) (**Figure 1**). Finally, there is an isoform-specific difference in the glutamate residue on the outer surface of the β-barrel facing the hydrophobic lipid core, near the middle of the lipid membrane: Glu73 (VDAC1) and Glu84 (VDAC2) (**Figure 1**). In contrast, VDAC3 has a corresponding glutamine (Gln73) at the same position.

**Figure 1.**
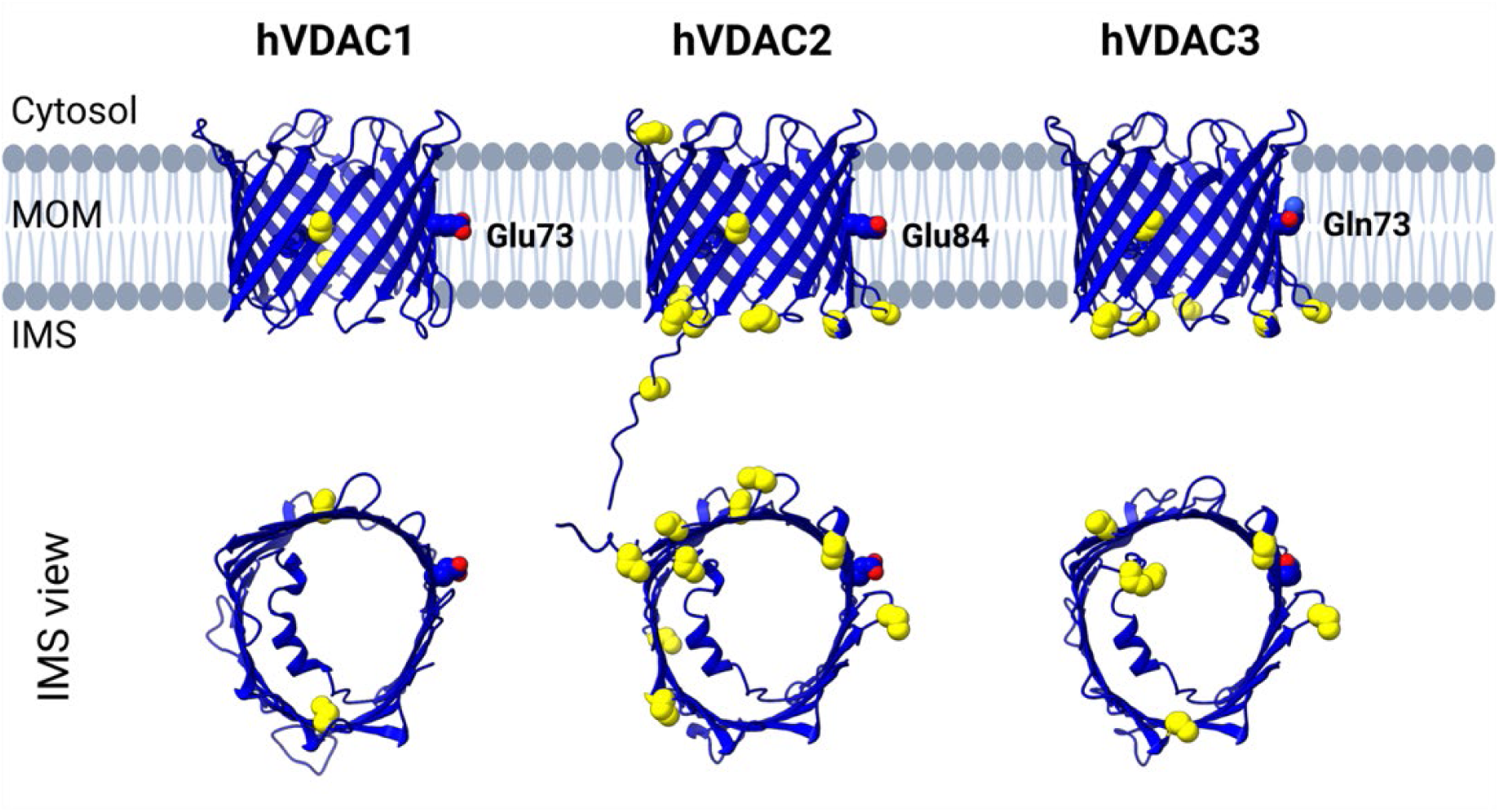
Structural comparison of human VDAC isoforms. Cartoon representation of three human VDAC isoforms. *Top*: VDAC1 (PDB: 6G6U) obtained from X-ray crystallography. Structures of VDAC2 and VDAC3 were generated based on sequence homology using ColabFold (99). *Bottom*: IMS view of VDAC1, VDAC2 and VDAC3. Cysteine residues are shown as yellow spheres to highlight the isoform-specific differences in cysteine numbers. Membrane-facing glutamate residues Glu73 (VDAC1) and Glu84 (VDAC2), and corresponding Gln73 (VDAC3), are labeled. Adapted with permission from (6).

When reconstituted into a planar lipid membrane (PLM) (**Figure 2A**), all three isoforms form almost identical, highly conductive channels which gate, i.e., transit from the unique high-conductance open state (∼ 4 nS in 1 M KCl) to a variety of low-conductance “closed” states in response to the applied voltage (**Figure 2B**). VDACs are known to have multiple voltage-induced closed states which are characterized by a wide range of conductances and low anionic or cationic selectivity (3, 49, 50). In contrast, the selectivity of open-state conductance is anionic with Cl^-^ to K^+^ permeability ratio *P_Cl-_/P_K+_* ∼ 1.3 (in 5x salt gradient of 1 M KCl *cis* / 2 M KCl *trans*), and similar for all three VDAC isoforms (51, 52) (**Figure 2C**). The voltage gating of reconstituted VDACs is conventionally analyzed using multichannel systems (53, 54). The normalized conductance (*G/G_max_*) plots for all three VDAC isoforms display the characteristic bell-shaped voltage dependences (**Figure 2D**). There is a measurable difference in the minimal conductance (the closed state conductance, *G_min_*), yet the open probability (*P_open_*) plots overlap for all isoforms (**Figure 2E**). These similarities indicate that they can all concurrently facilitate transport of the negatively charged metabolites, such as ATP and ADP, in and out of mitochondria. It is not surprising that the minor structural differences among the three VDAC isoforms do not affect their basic channel properties, such as conductance, ion selectivity, and voltage-gating. However, it remains a mystery how isoforms with similar structures and channel properties can exhibit specific cellular functions shown in cellular model systems and isoform-specific knock-out mice, where each VDAC isoform has distinctive functional roles (12, 15, 42, 43).

**Figure 2.**
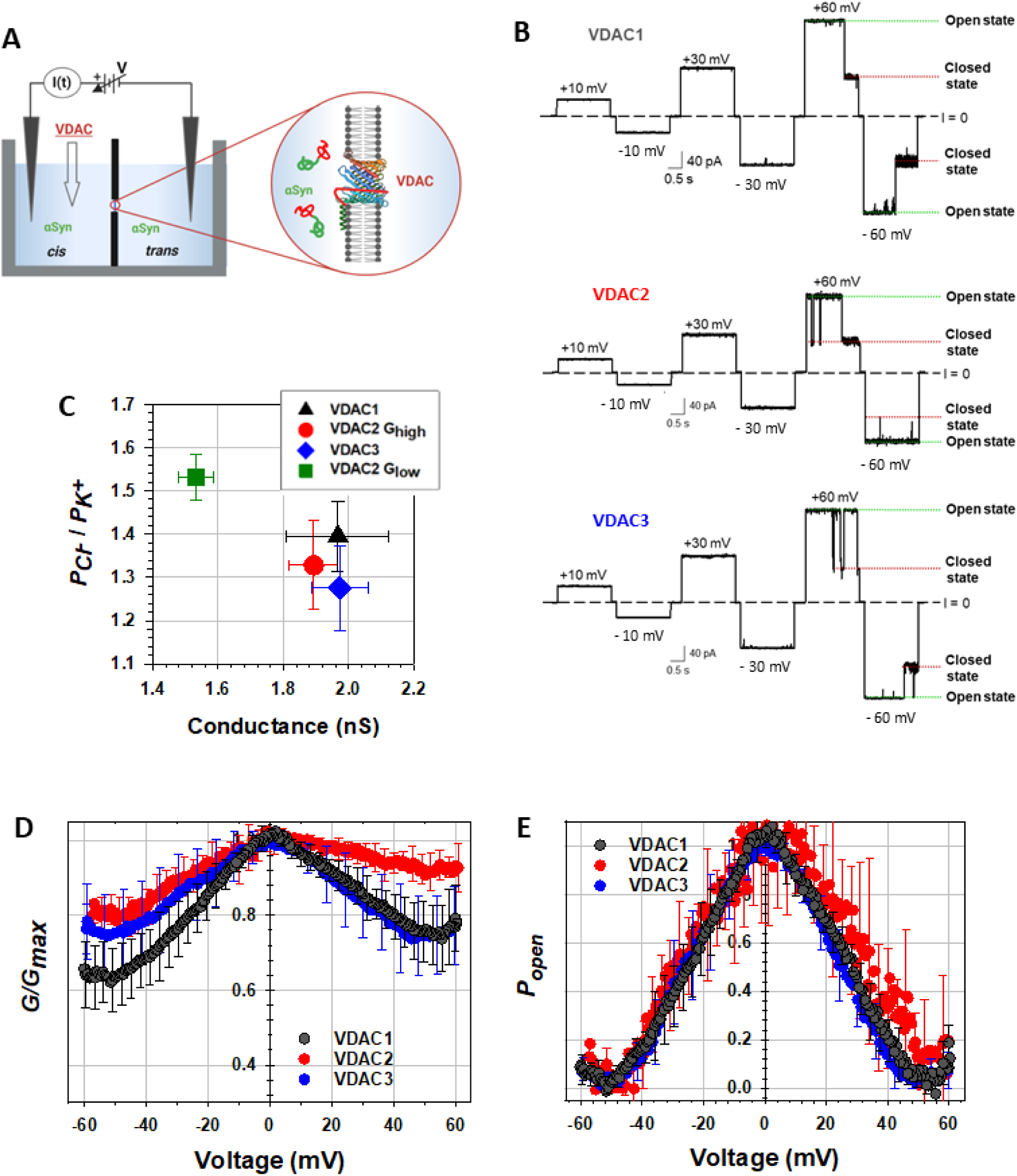
All three VDAC isoforms form similar high-conductance, anion-selective, voltage-gating channels when reconstituted into PLM. **(A)** A schematic of the experimental setup for VDAC reconstitution. In these experiments, VDAC is inserted into PLM when added to the “cis” compartment of the experimental chamber filled with buffer solution. The ionic current through the channel is generated by the applied voltage and constantly recorded in voltage-clamp mode. αSyn is added to both compartments. **(B)** Representative single-channel traces of wild-type mVDAC1, hVDAC2, and hVDAC3 reconstituted into PLM formed from 2DOPG/DOPC/DOPE lipid mixture in 1 M KCl (pH 7.4) at increasing voltages. Here, and elsewhere, dashed lines indicate the zero-current level; dotted lines indicate the open (green) and closed (red) states of VDAC. The current record was digitally filtered at 500 Hz using a low-pass Bessel (8-pole) filter. Modified with permission from (51, 52). (C) All three VDAC wild-type isoforms form anion-selective channels with similar permeability ratios P_Cl-_ /P_K+_ ∼ 1.3 for the fully open conductance states of 1.6-2.0 nS in 1.0 M (cis)/0.2 M (trans) KCl gradient (corresponding to a conductance of ∼4.0 nS in symmetrical 1.0 KCl). The exception is VDAC2 substates with conductance < 1.6 nS (VDAC2 G_low_) and high anion selectivity (P_Cl-_ /P_K+_ ∼ 1.5). Error bars are ±SD from at least five independent channel measurements for each VDAC isoform. PLM were formed from 2DOPG/DOPC/DOPE lipid mixture. Modified with permission from (51). **(D-E)** Characteristic bell-shaped plots of normalized average conductance *G/G_max_* (D) and open probability (*P_open_*) (E), as functions of the applied voltage for mVDAC1, hVDAC2, and hVDAC3. *G* is conductance at a given voltage, and *G_max_* is the maximum conductance at |V| ≤10 mV. *P_open_* was calculated as described in (52). Membranes were made of Polar Lipid Extract (PLE) in 1 M KCl (pH 7.4). Data are means of 3-11 independent experiments ± SD (error bars shown every 5 points for clarity).

Therefore, the natural question is what distinguishes the unique physiological function of these almost identical proteins? And what is the biophysical basis of such functions? We hypothesize that VDAC can regulate mitochondrial function through its interactions with multiple cytoplasmic and MOM proteins; isoform-specific complexation is the key to distinguishing between isoforms’ cellular functions (see our recent review (6)). Here, we focus exclusively on the biophysical aspects of distinct interactions between VDAC isoforms reconstituted into PLM with monomeric α-Synuclein (αSyn), using this intrinsically disordered cytosolic protein as a sensitive molecular probe.

## 2. α-Synuclein-VDAC interaction

### α-Synuclein

αSyn is a small, 140-residue, 14.4 kDa intrinsically disordered neuronal protein and is an established hallmark of Parkinson’s disease (PD), where αSyn in its fibrillar form constitutes a major component of Lewy bodies (55, 56). αSyn is primarily found in neural tissue, making up as much as 1% of all proteins in the cytosol of brain cells (57). It is predominantly expressed in the neocortex, hippocampus, substantia nigra, thalamus, and cerebellum. Though primarily a neuronal protein, it can also be found in peripheral tissues (58). Being mostly a cytosolic protein, αSyn was found to be associated with mitochondrial membranes preferentially over other native cell membranes (59–62). The increased level of αSyn expression is associated with its accumulation in fibrillar and oligomeric forms in dopaminergic neurons in PD (55, 56). Therefore, αSyn toxicity is conventionally related to its aggregates. However, the detailed mechanisms of αSyn involvement in neurodegeneration, particularly in its monomeric form, remain largely unknown.

Importantly, αSyn was found to be associated with both mitochondrial membranes – MOM and IM – causing mitochondrial dysfunction (59, 62–67). Overexpression of αSyn in SH-SY5Y cells leads to increased cell death and loss of mitochondrial membrane potential (**Figure 3A,B**) (68). Mitochondrial localization of αSyn was confirmed in HeLa (**Figure 3C**) and embryonic hypothalamic neuronal cell line (**Figure 3D**) using a proximity ligation assay (PLA) between αSyn and COX IV in the IM (69). Co-localization of αSyn with VDAC has been reported by several groups (reviewed in (70)). Furthermore, VDAC1 knockdown in neuroblastoma cells reduced αSyn proximity to COX IV, thus supporting VDAC1 as a pathway for αSyn across the MOM (68). We recently proposed that one of the physiological functions of monomeric αSyn is through its dynamic interaction with VDAC (24, 68). We demonstrated that the mechanism of αSyn-VDAC interaction could be successfully studied by using the electrophysiology approach on VDAC reconstituted in PLM.

**Figure 3.**
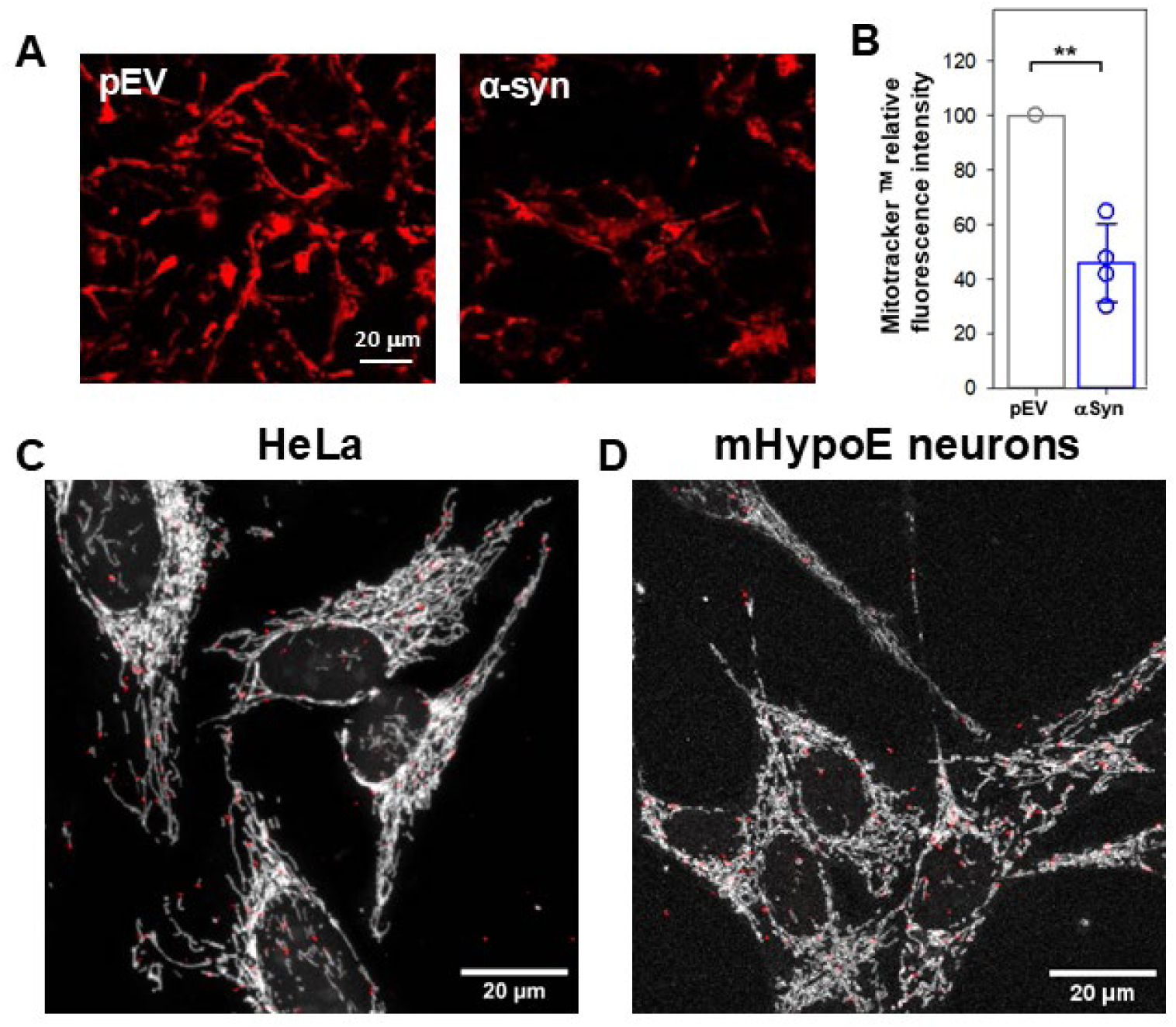
αSyn enters mitochondria, inducing mitochondrial dysfunction. **(A)** αSyn depolarizes mitochondria in dopaminergic cells. Representative images of differentiated SHSY5Y cells transiently transfected with empty vector (pEV) or WT αSyn. Cells are loaded with MitoTracker CMXRos to evaluate mitochondrial membrane potential (scale bar 20 μm). (**B**) The relative fluorescence intensity changes are quantified as ratios over pEV control presented as mean ±SD (n = 4). Significance was measured using one-way ANOVA (**p < 0.01). Modified with permission from (68). (**C**) Representative image of HeLa cells shows the αSyn and COX IV (red dots) colocalizing at IMS using Proximity Ligation Assay (PLA). Modified with permission from (69). (**D**) Representative image of PLA between αSyn and COX IV (red dots) colocalizing at IMS in embryonic mouse hypothalamus neuronal cell line (mHypo43). MOM is labeled with Tom20-Alexafluor488 (gray) (scale bar 20 μm).

### α-Synuclein interaction with VDAC

Using the method of ion channel reconstitution in PLM (**Figure 2A**), we showed that monomeric αSyn at nanomolar concentrations reversibly and partially blocks conductance of all three VDAC isoforms (24, 51, 52) (**Figure 4A**). αSyn has two domains: the membrane-binding domain and the negatively charged C-terminal tail (CTT) (**Figure 4B**). According to the earlier proposed model of αSyn-VDAC complexation, the first step is αSyn binding to the membrane by its N-terminal domain, followed by capture of its CTT by the VDAC pore (**Figure 4C**) (24). The capture rate, or the on-rate, depends exponentially on the applied potential (**Figure 4D**). Moreover, the CTT can retract back after some time spent inside the pore, which, in turn, depends on the magnitude of the applied voltage. This model includes only two functionally important steps of interactions – the αSyn binding to the membrane surface – the first step - and capture of the highly negatively charged CTT of αSyn by the VDAC pore driven by the electric field – the second step (71) (**Figure 4C**). Both steps are intricate in their own, e.g., as we have shown previously, the strength of the binding, defining the sheer amount of membrane-bound αSyn, is not a full descriptor in what relates to the on-rate of αSyn capture by the VDAC pore (72). The on-rate is strongly influenced by the conformation of the membrane-bound αSyn, specifically, by the availability of its CTT for capture. This conformation depends on the lipid species used for membrane formation (72) and can change with the surface concentration of the bound αSyn (73). The strong dependence of the on-rate on the efficiency of αSyn-membrane binding, and consequently, on membrane lipid composition, has been shown experimentally in single-channel experiments (72) for αSyn interaction with mouse VDAC1, where *k_on_* of αSyn interaction with VDAC1 reconstituted into the PLM made of anionic lipids (dioleoylphosphatidylglycerol (DOPG)/dioleoyl-phosphatidylcholine (DOPC)/dioleoyl-phosphatidylethanolamine (DOPE) mixture in a ratio of 2:1:1 (wt/wt), 2DOPG/DOPC/DOPE) was ∼100 times higher than in a positively charged PLM (using a positively charged dioleoyl-trimethylammonium-propane (DOTAP) mixture with DOPC and DOPE, 2DOTAP/DOPC/DOPE) (72). Many previous studies have shown that αSyn-membrane binding strongly depends on membrane lipid composition, with preferential binding to anionic lipids (74–76). The proposed mechanism has hydrophobic and electrostatic components of the absorption of αSyn N-terminal α-helix into the lipid headgroup region (76–78), which also explains the preference of αSyn for anionic and PE lipids (79). Therefore, it is instructive to compare the results of the single-channel experiments obtained in PLMs made of the same lipid composition.

**Figure 4.**
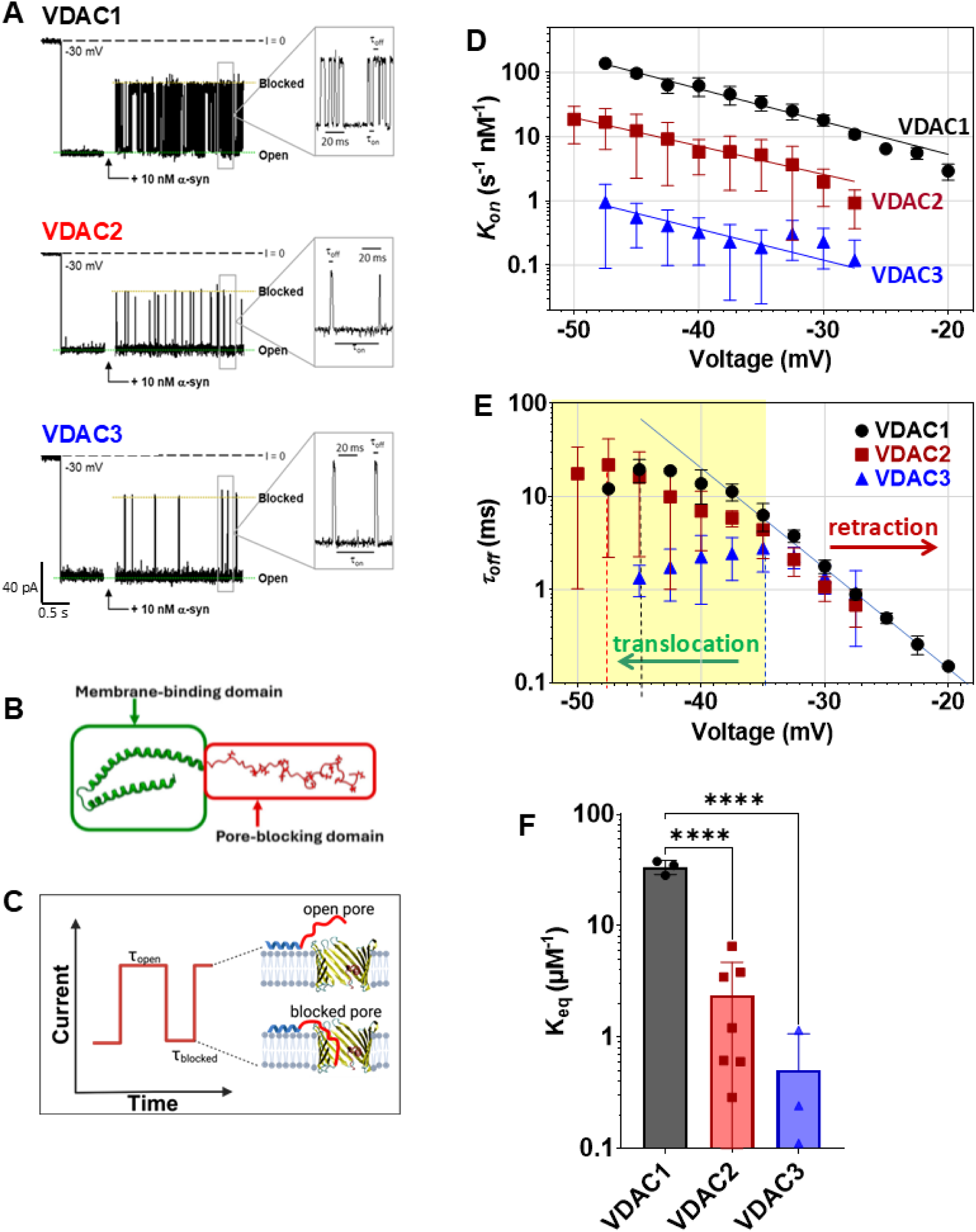
Kinetic interaction with αSyn distinguishes the three VDAC isoforms. (**A**) Representative mVDAC1, hVDAC2, and hVDAC3 single-channel current traces before (*left traces*) and after addition of 10 nM αSyn (*right traces*) at -30 mV of applied voltage. Insets show fragments of current records at a finer time scale. Current records were digitally filtered at 1 kHz using a low-pass Bessel (8-pole) filter. (**B**) Monomeric αSyn consisted of the membrane-binding N-terminal domain (in green) that prevents translocation of the whole molecule through the pore only up to a certain voltage, and the pore-blocking negatively charged C-terminal domain (in red). Adapted with permission from (6). **(C)** A model of αSyn-VDAC interaction. The blockage events are characterized by the time when the channel is open (*τ_on_*) and the time when CTT of αSyn is captured inside the pore (*τ_blocked_*), and the channel conductance is blocked. Adapted with permission from (71). (**D**) Voltage dependences of the on-rate constant *k_on_* = 1/(<τ_on_>[C]), of αSyn binding to mVDAC1, hVDAC2, and hVDAC3, where [C] is the αSyn bulk concentration. The solid lines represent a fit to the Boltzmann equation *k_on_* (V) = *k_0_* exp [*n_b_e|V|/(k_B_T)*], where *V* is the applied voltage, *n_b_* is the effective binding charge of 2.8 ± 0.2, and *e*, *k_B_*, and *T* have their usual meaning of elementary charge, Boltzmann constant, and absolute temperature, respectively. (**E**) Voltage dependences of the mean blockage time *τ_off_ = <τ_blocked_>* obtained for mVDAC1, hVDAC2, and hVDAC3. The increase of *τ_off_* with voltage amplitude corresponds to the αSyn blockage/retraction regime; the decrease of *τ_off_* with voltage amplitude (highlighted in yellow for VDAC3) corresponds to the regime of αSyn translocation through the pore. The solid line is to guide the eye. The vertical dashed lines indicate the voltage (*V*)* separating the two regimes for each isoform. Data are means of 3-8 experiments ± SD. Membranes were formed from mixtures of 2DOPG/ DOPC/DOPE/, and membrane bathing solutions contained 1 M KCl buffered with 5 mM HEPES at pH 7.4. **(F)** The equilibrium constant, *K_eq_*, for WT mVDAC1, hVDAC2, and hVDAC3. *K_eq_* = *k_on_*·τ_off_ was determined from the *k_on_* and *τ_off_* values obtained at -30 mV, presented in (D, E). Data are the means of 3-7 experiments ± SD. The significance was measured using an ordinary one-way ANOVA test (****p<0.0001).

We compared channel properties of all three VDAC isoforms reconstituted into PLM formed from 2DOPG/DOPC/DOPE lipid mixture, which mimics the high content of PC and PE lipids, with DOPG mimicking the negatively charged anionic lipids of the MOM (80). The single-channel recordings in **Figure 4A** show examples of similar conductance fluctuations for all three VDAC isoforms between the open and blocked states (with the latter being ∼0.4 of the open-state conductance), in the presence of 10 nM αSyn in the membrane-bathing 1 M KCl solutions in the membranes made from a 2DOPG/DOPC/DOPE lipid mixture. αSyn induces similar blockage events from both sides of the channel when a negative voltage is applied to the side of αSyn addition. For the sake of clarity, here and elsewhere in this work, we show single-channel current records and the experimental data at the negative potentials only. The αSyn-VDAC interactions obtained at both voltage polarities are shown in **Supplemental Figures S1 and S2**. In each VDAC isoform, αSyn induces characteristic, well-time-resolved blockage events on the time scale of 0.1-100 ms, which are qualitatively similar but quantitatively significantly different.

### Comparison of VDAC isoforms in their kinetic interaction with αSyn

While αSyn induces similar characteristic conductance blockages (millisecond time scale) in each isoform (**Figure 4A**), there is up to 100-fold difference in on-rates between VDAC1 and VDAC3, with the on-rate for VDAC2 in between, which is a remarkable indication of isoform specificity (**Figure 4D**) (24, 51, 52). The striking difference in the on-rate constant, *k_on_* (the inverse of <τ_on_> normalized over the αSyn concentration), between isoforms is observed for all voltages and at both polarities (**Supplemental Figure S1B**). Notably, the slopes of *k_on_* voltage dependences are similar for all isoforms, indicative of a similar effective “gating charge” of 3.0 ± 0.7. There is also some asymmetry in the voltage dependences of *k_on_* with respect to the polarity of the applied voltages (**Supplemental Figure S1B**), with the highest difference between the *k_on_* at negative polarity, but with the same sequence for *k_on_* decrease between isoforms: VDAC1>VDAC2>VDAC3. Thus, the pronounced difference in the on-rates of αSyn-VDAC interaction between isoforms cannot be merely assigned to the differences in αSyn membrane binding and therefore should involve the specificity of αSyn interaction with each isoform.

The increasing negative voltage keeps the anionic CTT of αSyn inside the VDAC pore longer, leading to an exponential increase of the blockage time, *τ_off_* (**Figure 4E**). At low applied voltages, the blockage time *τ_off_* increases in a voltage-dependent manner and remains quite similar for all three isoforms (**Figure 4E**, solid line). Under low applied voltage, αSyn is captured by the pore and then retracts back to the same side of the channel (**Figure 4C**) (81). This is a retraction regime. The release rate becomes more complex at higher voltages, which is seen as biphasic behavior of the blockage time *τ_off_* with voltage, indicating the onset of αSyn translocation through VDAC (**Figure 4E**, yellow highlight). Importantly, the applied voltage is the external force that pulls the blocking domain into the VDAC pore and, upon exceeding some threshold (usually > ׀30׀ - ׀40׀ mV, depending on membrane lipid composition, salt concentration, and VDAC isoform type), drives the whole αSyn molecule to the opposite side of the membrane. Further experimental confirmation of this model came from single-channel selectivity experiments, where the selectivity of the αSyn-blocked state of VDAC was found to be reversed to cationic due to the presence of highly anionic CTT inside the pore (82, 83). The mechanism of biphasic behavior of *τ_off,_* indicating the voltage-induced translocation, was previously described for the blockage of anthrax toxin channels by charged molecules (84).

The voltage amplitude (*V\**) required to induce the translocation regime varies quite remarkably from -35 mV for VDAC3 to -45 mV for VDAC1 (indicated by vertical dashed lines in **Figure 4E**), implying their different ability to translocate αSyn into mitochondria. The data for VDAC2 also show off-rate saturation, but do not allow for a reliable evaluation of *V\** because of the high spread of the datapoints. The similarity of *τ_off_* and the difference in *k_on_* for VDACs WT isoforms (**Figure 4D,E**) yield a pronounced difference – up to 100 times – in the equilibrium constant, *K_eq_* = *k_on_* * *τ_off_*, of the αSyn-VDAC interaction between three VDAC isoforms (**Figure 4F**). There is more than an order-of-magnitude difference in the equilibrium constants between WT VDAC isoforms (**Figure 4F**). This is despite their essentially identical open-state conductance and ion selectivity (**Figure 2B,C**).

We can speculate that while VDAC3 expression level in most cells is significantly lower than that of VDAC1 and VDAC2, the blockage on-rate of VDAC3 by αSyn is ∼100 times lower, which could result in VDAC3 being mostly open for ATP/ADP transport while VDAC1 is blocked by αSyn, thus making VDAC3 the most important VDAC isoform for respiration (35, 85). On the other hand, ∼10 mV lower voltage is required to induce αSyn translocation through VDAC3 compared with VDAC1, further emphasizing the specific role of each isoform in respiration and in αSyn-induced mitochondrial toxicity.

### The effect of cysteines on αSyn-VDAC interaction kinetics

One of the differences between VDAC isoforms is the number and position of cysteine residues (86) (**Figure 1**). VDAC1 and VDAC2 share one cysteine outside of the β-barrel facing the lipid core – Cys127 and Cys138, respectively. Another residue, Cys232, unique to VDAC1, is positioned inside the pore close to the IMS side. The other six VDAC2 cysteine residues are located on the loops, with one facing the cytosol and five on the IMS-facing loops. VDAC3 has four cysteines exposed to the IMS (87). Both VDAC2 and VDAC3 also have two cysteines on the N-terminal α-helix (87). There is a notable asymmetry in cysteine distribution in these isoforms between the loops facing the cytosolic and IMS sides, with most cysteines crowded at the IMS interface, leading De Pinto and coauthors to propose that four IMS-facing cysteine residues in VDAC3 could be the redox sensors important for channel function (88, 89). VDAC2 has more cysteines, but research has focused on VDAC3’s role in ROS production and the role of its cysteines (86, 88, 90, 91). Future systematic research on the physiological role of VDAC2 cysteines is needed.

We hypothesized that cysteine residues may affect VDAC interaction with αSyn and thus could be the source of the observed difference between isoforms. We tested this hypothesis by comparing the kinetics of αSyn interaction with VDAC1, 2, and 3 cysteine-less mutants (ΔCys) with their corresponding WT isoforms. We have previously shown that the average open-channel conductance for VDAC3-ΔCys and VDAC1-ΔCys had no measurable changes in comparison with WT, and the channels maintained their mild anionic selectivity (52) (**Supplemental Figure S3A-D**). The situation with VDAC2 conductance is in general more complex, because VDAC2, as we have recently demonstrated, stochastically switches between multiple high-conductance states, which results in a wide conductance distribution of the open state in the range of 2.9 to 4.1 nS (in 1 M KCl) (**Supplemental Figure S3E,F**), uncommon for other VDACs (51). Notably, the selectivity of these high-conducting substates of VDAC2 WT remains anionic (**Figure 2C**); therefore, they should be distinguished from the well-described low-conducting (closed) voltage-induced states, which are characterized by cationic or less anionic selectivity (51). VDAC2-ΔCys mutant, reconstituted into the same 2DOPG/DOPC/DOPE membrane, forms channels with 3.4-4.4 nS conductance as VDAC2 WT, with the transitions to the lower conductance substates (< 3.8 nS) observed only in 20% of experiments (in comparison with 50% of VDAC2 WT (51). Therefore, here we compare the data obtained for VDAC2-ΔCys channels of 3.8-4.0 nS conductance with the average data for all conductance substates obtained for VDAC2 WT. This explains the high standard deviations obtained for VDAC2 samples.

We found that while cysteine residues do not have a measurable effect on VDAC isoforms’ conductance and ion selectivity, they affect VDAC affinity for αSyn, especially for VDAC3 (52). The systematic comparison of the voltage dependences of *k_on_* between the ΔCys mutants and the WT for each isoform is shown in **Figure 5A,C,E**. At negative applied voltages, the on-rate of αSyn-VDAC3-ΔCys interaction is ∼ 10 times higher than with VDAC3 WT (**Figure 5E**). For VDAC2-ΔCys, the *k_on_* is ∼5 times systematically higher than for VDAC2 WT, but with low significance (95% confidence interval for exponential dependence at ׀*V*׀ > 35 mV) (**Figure 5C**). In contrast, there is no significant difference between *k_on_* values for VDAC1-ΔCys and the WT (**Figure 5A**). At positive voltages, the ∼3-fold increase in *k_on_* for VDAC2-ΔCys is comparable with the increase at negative voltages (**Supplemental Figure S2**). However, the *k_on_* at positive voltages for VDAC3-ΔCys and WT are essentially the same (**Supplemental Figure S2**). There is no significant difference in *k_on_* values between VDAC1-ΔCys and WT for both polarities of the applied voltage (**Supplemental Figure S2).** Importantly, the slope of the *k_on_* voltage dependence (solid lines in **Figure 5A,C,E**) does not change with cysteine removal for all isoforms, WT and cysteine-less, leading to about 3.1± 0.6 effective “binding charge” for all VDAC samples.

**Figure 5.**
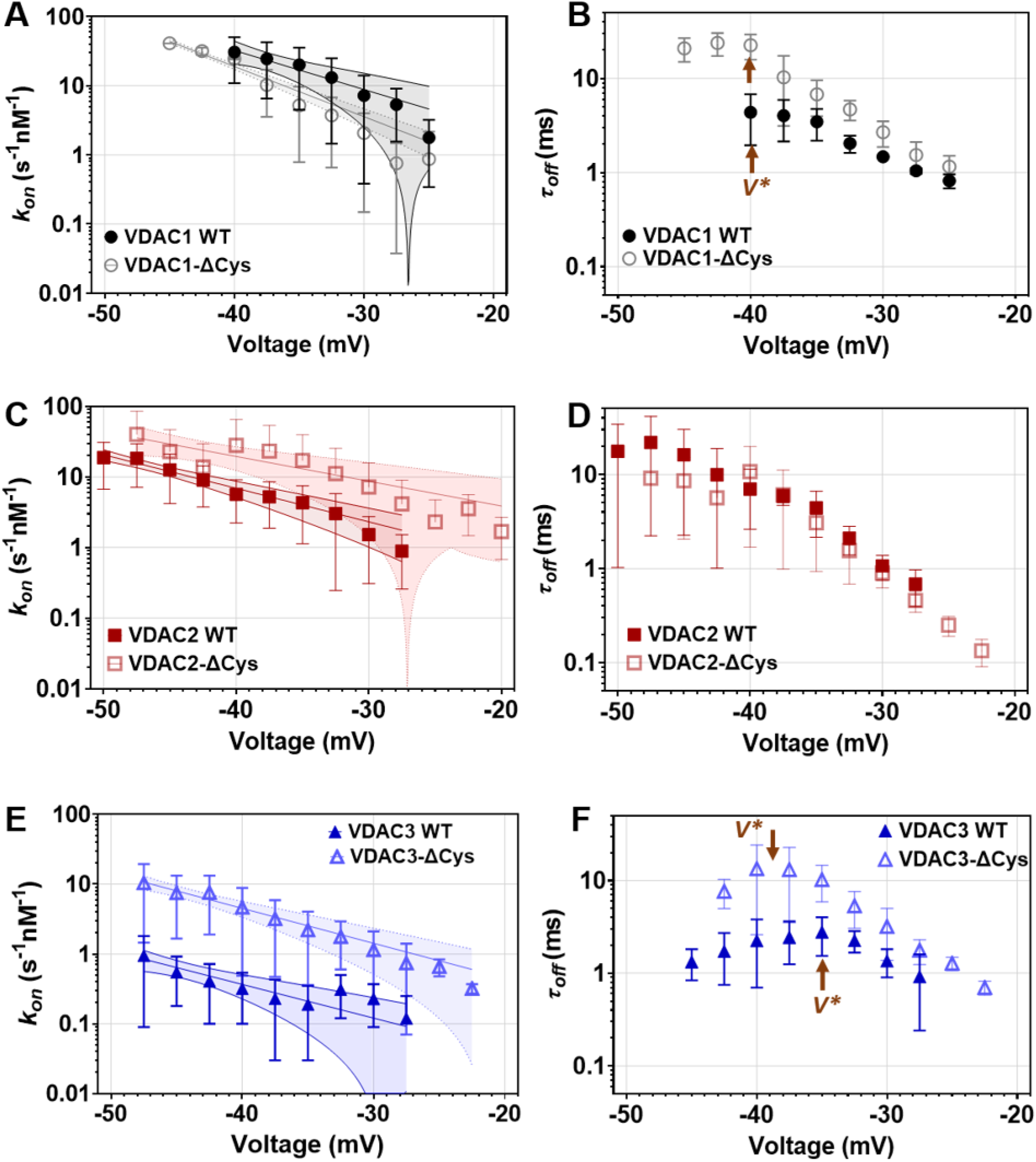
Cysteines differently affect the kinetics of αSyn interaction with VDAC isoforms. Voltage dependences of the on-rate constants *k_on_* (**A, C, E**) and the mean blockage time τ*_off_* (**B, D, F**) of αSyn binding to hVDAC1 and hVDAC1-ΔCys mutant (**A,B**), to hVDAC2 and hVDAC2-ΔCys mutant (**C, D**), and to hVDAC3 and hVDAC3-ΔCys mutant (**E, F**). The solid lines in (A, C, E) represent a fit to the Boltzmann equation (as in the Figure 4 legend), fitted with the effective “binding charge” *n_b_* varied from 2.1 to 4.3 (<*n_b_* > = 3.1± 0.7) between VDACs. Shading shows the 95% confidence band of the fit calculated in GraphPad Prism version 11.1.0. *V\** indicates the voltage separating the blockage/retraction and translocation regimes. Data are the means of 3-7 independent experiments ± SD. Membranes were formed from 2DOPG/DOPC/DOPE/ lipid mixtures in 1 M KCl buffered with 5 mM HEPES at pH 7.4

The blockage time in the retraction regime, characterized by exponential rise of *τ_off_* with absolute voltage magnitude, is not significantly affected by cysteine removal for all isoforms (**Figure 5B, D, F**). The most pronounced difference in *τ_off_* between ΔCys and WT VDACs was found for VDAC3: the voltage *V\** at which the retraction regime switches to the translocation regime is 5-10 mV higher for VDAC3-ΔCys than for the WT (**Figure 5F**), indicating that cysteines in VDAC3 may facilitate αSyn translocation through the pore. This effect takes place at both voltage polarities for VDAC3-ΔCys (**Supplemental Figure S2**). To conclude, the time αSyn spends inside VDAC pore is not affected by cysteine removal, with only VDAC3 being a possible exemption.

*K_eq_* values for VDAC2 and VDAC3 cysteine-less mutants are also ∼3-6 times systematically higher than for the WT VDAC2 and VDAC3 (significant for VDAC3 with *p <0.05, but not significant for VDAC2 with p=0.2 in unpaired t-test with Welch’s correction) and essentially similar for VDAC1-ΔCys and the WT (*p=0.8) (**Figure 6**).

**Figure 6.**
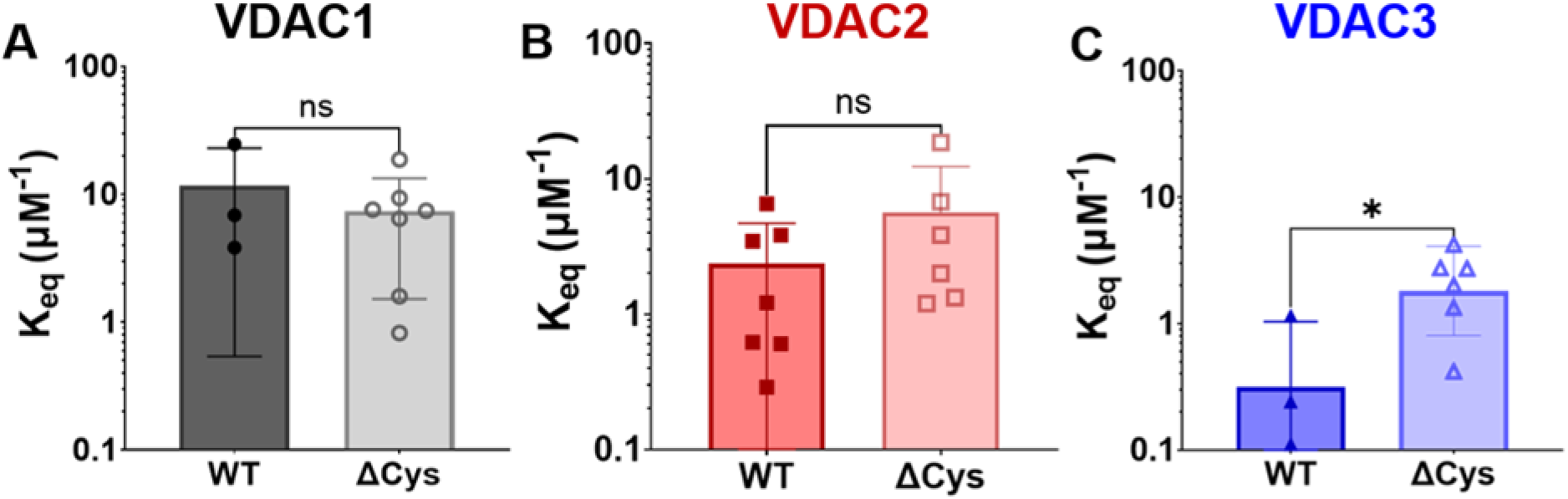
Cysteines affect αSyn interaction with the VDAC3 isoform leading to markedly different equilibrium constants. A comparison of *K_eq_* values between hVDAC1 (**A**), hVDAC2 (**B**), and hVDAC3 (**C**) WT and their corresponding ΔCys mutants. *K_eq_* = *k_on_*·τ_off_ were determined from the *k_on_* and *τ_off_* values obtained at -30 mV, presented in Figure 5. Data are the means of 3-7 experiments ± SD. The significance was measured using the unpaired t-test with Welch’s correction (*p<0.05).

The pronounced differences in equilibrium constants between three VDAC WT isoforms and between VDAC2 and VDAC3 WT and their cysteine-less mutants appear to be in striking contrast with the preserved *K_eq_* found for different VDAC2 conducting substates (**Figure 7**). There is a remarkable correlation between the conductance of VDAC2 open-state substate and the on- and off-rates of αSyn-VDAC2 interaction: as conductance of the substate increases, *k_on_* increases but *τ_off_* decreases, resulting in an essentially constant *K_eq_* (**Figure 7B**) (51). Similar relations between VDAC open state conductance and *k_on_* and *τ_off_* were also found for two VDAC2 mutants – hVDAC2 with truncated N-terminus first 11 residues, ΔN-VDAC2, and VDAC2 E84A mutant (51). Both mutants also showed a positive correlation between conductance and *k_on_* and a negative correlation between conductance and *τ_off_*, resulting in an essentially constant *K_eq_* between conductance levels for each mutant (51). We explained these intriguing results using the formalism of the transition path energy landscapes for VDAC2 substates (51). We concluded that the regulation of all VDAC2 substates by αSyn is described by the same affinity, suggesting that when αSyn molecule is captured inside the pore, its physical state and interactions with the pore are the same for all substates but differ substantially in its kinetic parameters (51). In contrast, the different *K_eq_* for different isoforms suggests that αSyn molecules interact differently with each VDAC. The effect of cysteines on *K_eq_* supports this model. The presence of multiple cysteines in VDAC2 and VDAC3 affects the interaction of αSyn with their pores; this is especially pronounced for VDAC3, for which even *τ_off_* is affected by the removal of cysteine residues. The voltage asymmetry in the on-rate of αSyn interaction with VDAC3-ΔCys fits with a highly asymmetric distribution of cysteine residues on opposite sides of the channel (**Figure 1**). Consequently, we can speculate that two cysteines conservatively located on the N-terminal α-helices of VDAC2 and VDAC3 are unlikely to play an essential role in interaction with αSyn molecule, because the main difference is in cysteine distribution on the loops of VDAC2 and VDAC3. These considerations allow us to suggest that cysteines located on the loops at the IMS side of the channel (**Figure 1**) are the most likely to be involved in interaction with αSyn, affecting the reaction on-rate. The absence of an effect of the removal of two cysteines in VDAC1 on the kinetics of its interaction with αSyn supports this conclusion. Interestingly, this seems to be a counterintuitive scenario, because it was natural to expect that the cysteines inside the pore should affect αSyn capture rate.

**Figure 7.**
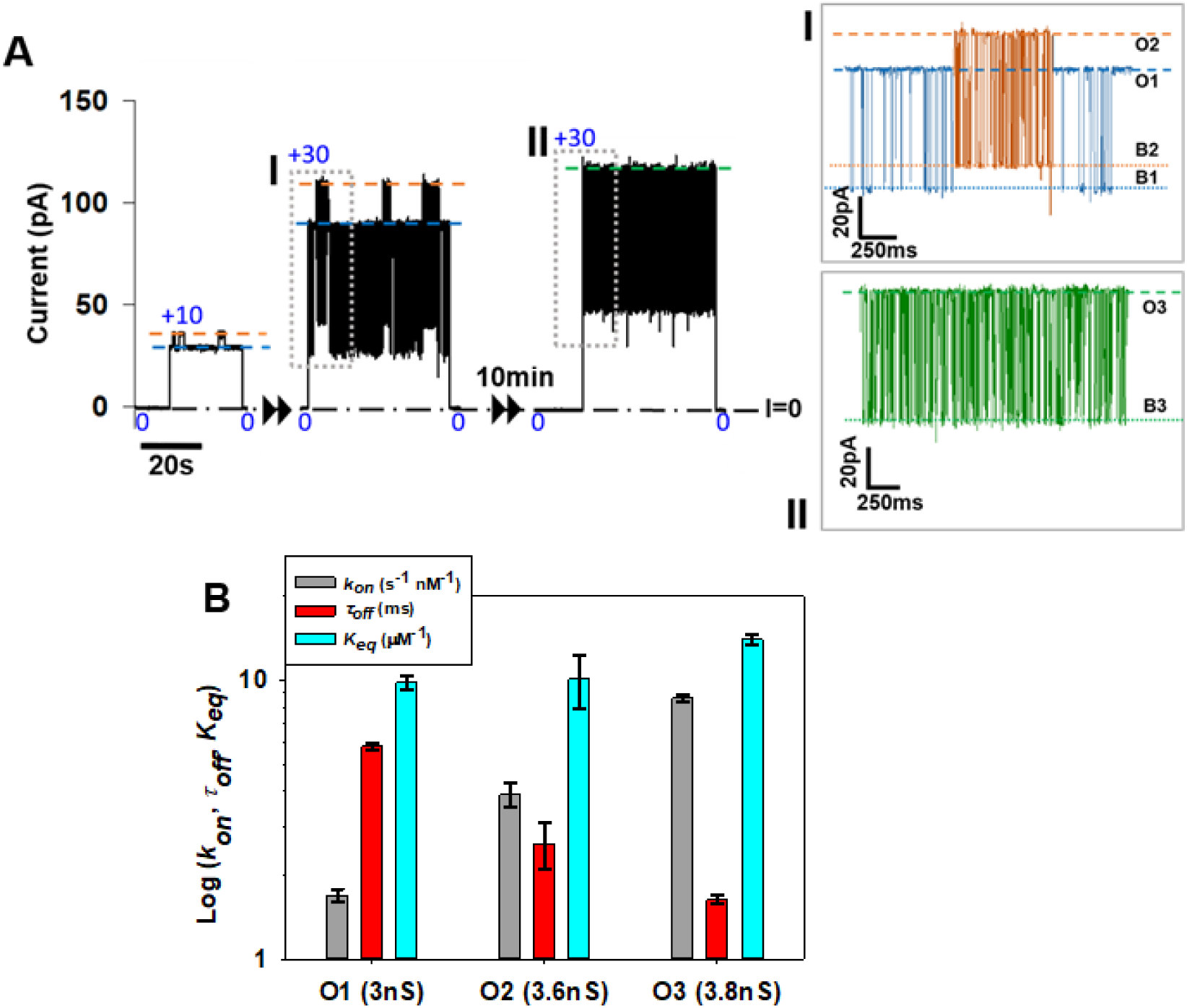
αSyn interacts with different high-conductance substates of VDAC2 WT with the same equilibrium constants. (**A**) A representative current record of a single hVDAC2 WT channel in the presence of 10 nM αSyn on both sides of the membrane formed from 2DOPG/DOPC/DOPE lipid mixture in 1 M KCl (pH 7.4) at 10 and 30 mV of applied voltages. The channel spontaneously fluctuates initially between two conductance substates of 3 and 3.6 nS indicated as O1 and O2, respectively, in Inset I (gray box). After ∼10 min of the continuous record, the channel spontaneously transitioned to the 3rd conductance state, O3, of 3.8 nS (Inset II) (gray box). The corresponding blocked states are shown in the Insets as B1, B2, and B3. (**B**) Comparison of the kinetic parameters of αSyn-VDAC2 interaction obtained in the experiment shown in (A). While the on-rate constant *k_on_* increases with an increase in the substate conductance and *τ_off_* decreases, respectively, the *K_eq_* remains constant. Adapted with permission from (51).

VDAC2 holds a special position among the three isoforms, being crucially important in development (92). However, the biophysical underpinning explaining its special role in cell physiology remains unexplored. We can only speculate that the unique plasticity of this channel and its interaction with αSyn (and other cytosolic partners) is key to understanding its uniqueness among VDAC isoforms. For instance, the preservation of equilibrium constants in interaction with αSyn was only found for VDAC2 different conductance substates, which points towards the specific nature of these substates.

VDAC2-αSyn on-rates fall between VDAC1 and VDAC3, depending on which conducting substate it exists in (51). We suggest that such interactive versatility is one of the important features of this unique isoform. The unexpected finding that substates vary in the kinetics of αSyn’s interaction with VDAC2 but not in its affinity implies that substates do not tune αSyn regulation of channel fluxes. However, the altered kinetics demonstrate that the capture of the synuclein molecule is affected by the structural changes in the channel.

Interestingly, a similar correlation between the channel conductance and the on-rate of αSyn capture was found upon comparison of three different β-barrel channels: α-hemolysin, MspA, and VDAC (93). It was shown that VDAC1, the channel with the highest conductance (4 nS in 1 M KCl) among the three channels, had ∼ 1000 higher on-rate than α-hemolysin, the channel with the lowest conductance of 0.73 nS in 1 M KCl. However, the equilibrium constant was not conserved for these three channels, similarly to what is reported here for the three VDAC isoforms with the same channel conductance.

## 3. Changes in free energy profile rationalize VDAC isoform differences

The two orders of magnitude difference in the αSyn capture on-rates, found for different VDAC isoforms reconstituted into PLM of the same lipid composition, can be rationalized by introducing an additional step in the capture reaction. The uncovered step of the capture process, reflecting direct interaction of αSyn with the membrane-VDAC interface, can be represented by an additional potential well along the reaction coordinate. **Figure 8**, which shows a tentative free energy profile, depicts it as state (**) located in between the state describing membrane surface-bound αSyn (*), and state (***), corresponding to the αSyn C-terminal tail residing in the pore. In the model, the interaction with the membrane-VDAC interface does not involve the C-terminal tail, as its contribution extends to state (***), thus increasing the relative depth of its potential well by the same *ΔU* (^3,1^). The change in the latter upon transition from VDAC3 to VDAC1 can be evaluated as 4 to 5 *k_B_T* units, which corresponds to about a hundred-fold change in the equilibrium constant. Importantly, this interaction cannot be explained by the effects of some small changes in isoforms’ conductance or selectivity (**Figure 2D**) because it was proven in our experiments with VDAC2 that its substates differing by 15-20 % by these parameters have shown equilibrium constants identical to that of the fully open state (**Figure 7B**).

**Figure 8.**
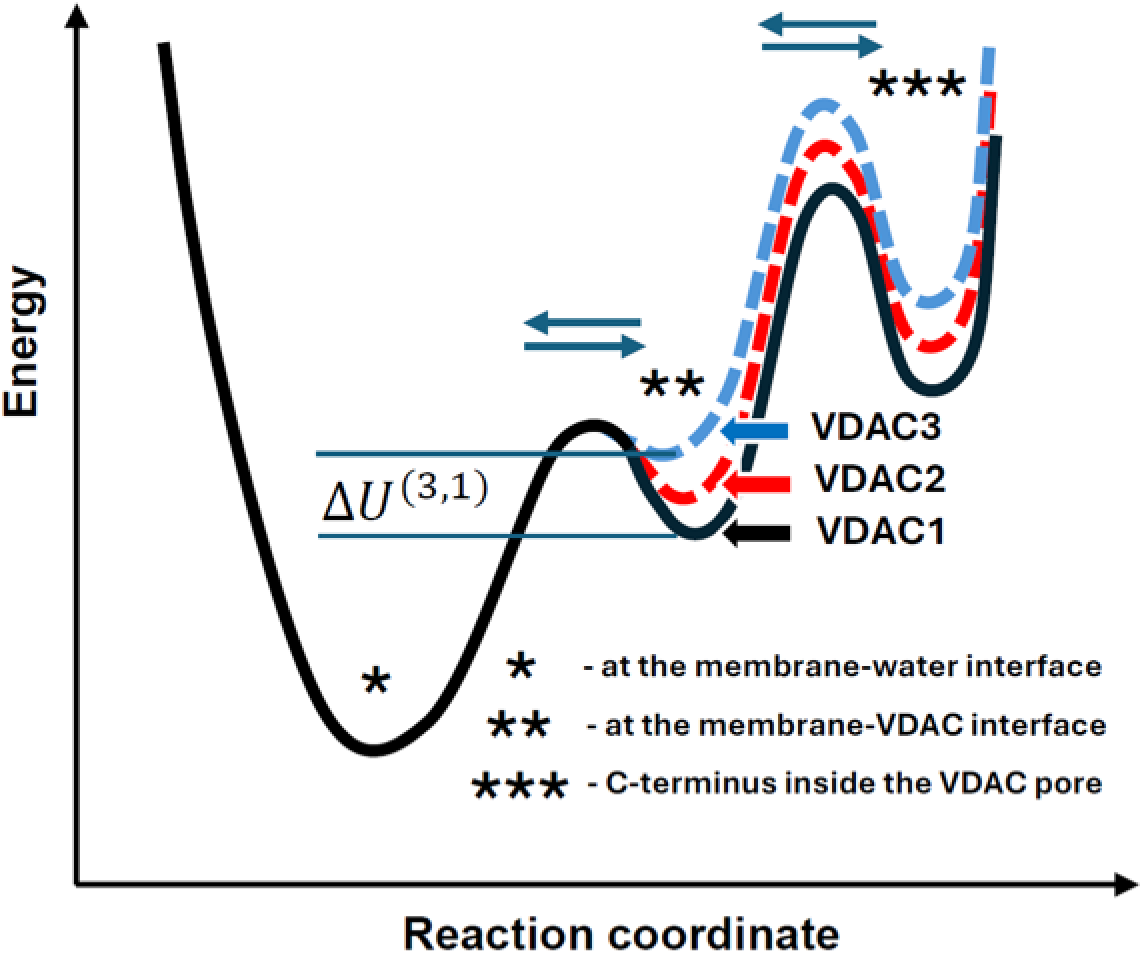
Proposed free-energy profile as a function of the reaction coordinate for the interaction of αSyn with VDAC isoforms, including a new important step involving the interaction of αSyn with the VDAC barrel–membrane interface. The concentration of αSyn in its membrane-bound state (*) can be differently enriched in the immediate vicinity of the channel through direct interaction with the VDAC barrel-membrane interface, corresponding to state (**). Among the three isoforms, the probability of finding αSyn in state (**) is greatest for VDAC1, resulting in the highest equilibrium binding constant for this isoform. The profile suggests that this occurs because the on-rate of the transition from state (**) to the state with the C-terminus inside the pore (***) is increased due to the higher occupancy of state (**) for VDAC1, while the residence time of the C-terminus inside the VDAC pore remains unchanged.

Such VDAC isoform-specific interaction is expected to change the characteristic voltage *V\** separating regimes of the C-terminal tail retraction and the voltage-driven whole molecule translocation for different isoforms. The increased interaction of the αSyn molecule with the membrane-VDAC1 interface is expected to shift *V\** to higher absolute values, as αSyn translocation would be hindered by such interaction. This is exactly what one can see in **Figure 4E** from the data displayed there. Indeed, the residence time curves suggest that compared to VDAC3, the onset of the translocation regime of αSyn through VDAC1 is delayed by about 10 mV. For the case of VDAC2, the transition voltage is undefined because of the significant spread of datapoints, but the value of *V\** in between those for VDAC3 and VDAC1 would be compatible with the data, considering the error bars given in **Figure 4E** for the VDAC2 datapoints.

Though solidly established in our experiments with VDAC isoforms, the mechanism of such direct interactions between αSyn and VDAC-membrane surface remains unknown and will be the goal of future computational and biophysical experiments on the identification of αSyn-VDAC binding sites, located, according to our model, in the vicinity of channel entrances that include the external loops of the VDAC structure. Meanwhile, an important piece of information that advances our understanding comes from the MD simulations of VDAC1 and VDAC2 isoforms in the membranes of the same lipid composition (2DOPG/DOPC/DOPE) that was used in single-channel experiments. A snapshot of the MD results is shown in **Figure 9**. It demonstrates a significant redistribution of lipids in the vicinity of VDAC. It is worth mentioning recent work that showed, using MD simulations, that VDAC1 influences the local lipid distribution within the membrane, creating distinct lipid layers (94). Most telling is our finding that redistribution of lipids is VDAC isoform-specific, meaning that, notwithstanding the high homology between the primary sequences of VDAC1 and VDAC2 and their practically identical structures (**Figure 1**), the outer surfaces of their β-barrels exhibit distinctly different interactions with membrane lipids. **Figure 9** shows that the effect of lipid-protein surface interaction is not only inhomogeneous along the channel outer circumference but is also drastically different for these VDAC isoforms. This immediately leads to two lines of reasoning. First, if interaction with lipids is so different, then it’s natural to expect similar, or even greater differences in the direct interactions with αSyn. Second, the VDAC isoform-specific significant distortion of the lipid composition around the channel means significant distortion in the lipid packing stress (95), leading to lipid packing defects in the close proximity to the channel. According to the literature, the latter have a profound effect on αSyn membrane binding (81, 96) and, therefore, may (partially) account for the effects under discussion.

**Figure 9.**
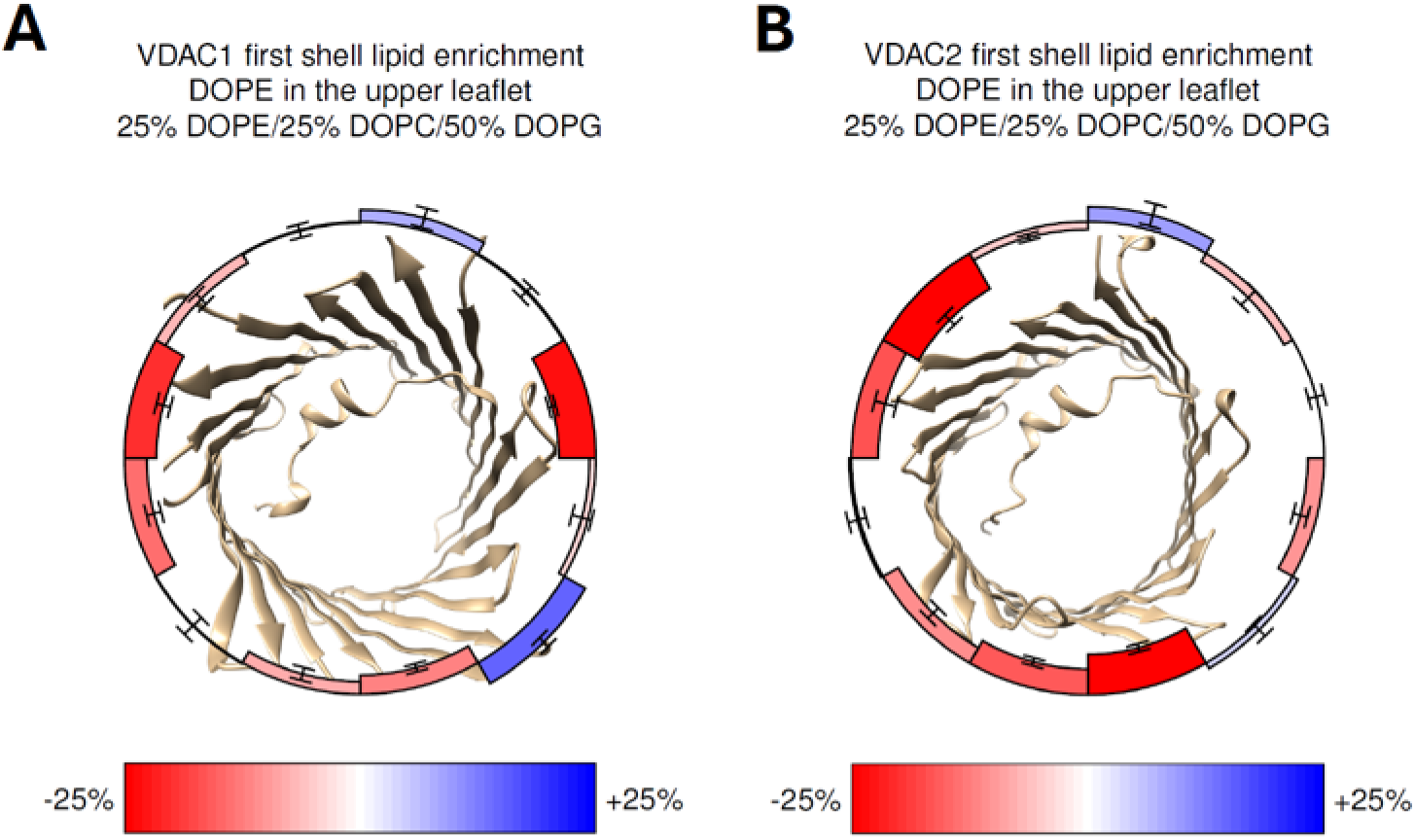
Molecular dynamics simulations demonstrate significant redistribution of lipids in the first shell of VDAC lipid environment of different VDAC isoforms. Both VDAC1 and VDAC2 enrich or deplete DOPE by as much as 25% in comparison to its average 25% molar concentration in the 2DOPG/DOPE/DOPC mixture. The effect is strongly dependent on the isoform, with VDAC2 accommodating more DOPE lipids at its outer surface. DOPE redistribution is also inhomogeneous along the surface circumference, with some structural fragments attracting DOPE lipids in excess of their average membrane concentration, and some depleting them.

Thus, a careful comparison of the changes in the dynamics of αSyn blockage between the three VDAC isoforms, which also takes into account recent results on the dynamics for the VDAC2 substates (51), brings crucial new insights into the interactions involved. It highlights the importance of the step that was absent in the original model, the step of direct αSyn interaction with the region of the membrane-VDAC interface. The importance of this step follows immediately from the magnitude of its effect, described by a 100-fold change in the equilibrium constant of the reaction, which recalculates into the change in the free energy of the blocked state by about 4 to 5 *k_B_T* units.

## 4. Physiological implications of the differential αSyn interaction with VDAC isoforms

The above biophysical results allow us to speculate that, on a cellular level, the less efficiently blocked VDAC3 remains open for respiratory substrate transport, while VDAC1 remains blocked by αSyn. Our recent comparative analysis of metabolism in HeLa cells with individual VDAC isoform KO supports this model: only VDAC3 KO decreases mitochondrial respiration, while VDAC1 KO affects glycolysis and VDAC2 KO does not measurably affect glycolysis or mitochondrial respiration (85). Similarly, VDAC3 was found to be critical for maintaining mitochondrial potential using VDAC isoforms knockdown in HepG2 cells (97). Since VDAC3 is blocked 10 times less effectively by dimeric tubulin than VDAC1 (52), we can anticipate that VDAC3 is primarily open while VDAC1 is closed via tubulin interaction. Thus, surprisingly, VDAC3, the least expressed isoform, turned out to be a critical one for mitochondrial function to fuel the high-energy demand of cancer cells.

Given that the mechanism of αSyn and tubulin interaction with VDAC is surprisingly similar, we believe that other undiscovered VDAC regulators could modulate metabolite fluxes and mitochondrial calcium through VDAC and hence permeability across MOM in an analogous fashion. The observed differential sensitivity of individual VDAC isoforms to αSyn and tubulin could also be a general pattern with other potential VDAC regulators, thus adding another physiologically important level of complexity to VDACs’ already multifunctional involvement in mitochondrial metabolism, calcium signaling and apoptosis. Thus, differences in VDAC regulation by cytosolic proteins may be a general mechanism defining the isoform-specific roles of VDACs *in vivo*. We can extrapolate this conclusion, stating that one of the distinguishing features of each isoform is its differential regulation by and interaction with other, yet unknown protein partners (6, 70, 98).

The question as to why VDAC2 KO is lethal or partially lethal, but KOs of the other two isoforms are not, remains open. We speculate that VDAC2’s unique plasticity, manifested by dynamic transitions between high conductance substates revealed in *in vitro* single-channel experiments, could be a key to its unique physiological significance. The same VDAC2 molecule could dynamically adapt its interaction with other proteins, similarly to what we found for the αSyn-VDAC2 complex. These data could tentatively explain the physiological significance of VDAC2: its ability to dynamically adjust to metabolic conditions and to change the rate of interaction with its presumably multiple protein partners.

Altogether, these results show that VDAC isoforms, being structurally and functionally similar—by the biophysical properties of conductance, selectivity, and voltage-gating—vary most drastically by their interactions with *α*Syn, thus suggesting that it is precisely the interaction with cytosolic regulators that clearly distinguishes each VDAC isoform and likely explains the origin of their biological variance.

Moreover, the presented findings point toward a new approach in our search for an effective drug for VDAC pharmacology. Specifically, future efforts should not be limited to finding the best binders to the outer mitochondrial membrane with the best bound conformations that make the anionic disordered C-terminus (or its synthetic analog) readily available for capture by the VDAC pore (96); it is now imperative to look for the optimization of the direct, VDAC isoform-specific step of the blockage reaction established in the present study. We believe that this line of research opens a new strategy in the search for an effective drug for VDAC pharmacology.

## Acknowledgements

Figures 1, 2A, and 4C were created with a licensed version of BioRender.com. Present addresses of the authors:

William M. Rosencrans, National Institute of Neurological Disorders and Stroke, National Institutes of Health, Bethesda, MD, USA.

María Queralt-Martín, Laboratory of Molecular Biophysics, Department of Physics, Universitat Jaume I, E-12071 Castellón, Spain.

Mya S. Wolfe, Wright State University Boonshoft School of Medicine, Dayton, OH, USA. Bethel G. Beyene, Department of Chemistry and Biochemistry, University of Maryland, Baltimore County, Baltimore, MD, USA.

Motahareh G. Larimi, National Institute of Diabetes and Digestive and Kidney Diseases, National Institutes of Health, Bethesda, MD, USA.

## Author Contributions

T.K.R. and S.M.B. conceived and designed research; T.K.R., W.M.R., M.Q.-M., M.S.W., and M.R. analyzed data; W.M.R., M.Q.-M., M.S.W., N.A.B. performed experiments; B.G.B. and M.G.L. purified protein; T.K.R., M.R., W.M.R., M.Q.-M., and S.M.B. interpreted results of experiments; A.J.S. performed MD simulations; S.M.B. performed theoretical modeling; T.K.R., M.R., A.J.S. and S.M.B. prepared figures; T.K.R. drafted manuscript; T.K.R., M.R., and S.M.B. edited and revised manuscript; T.K.R., M.R., W.M.R., M.Q.-M., M.S.W., N.A.B., B.G.B., M.G.L., A.J.S., and S.M.B. approved final version.

