## Supplemental Material for "So similar, yet so different: α-synuclein recognition of the isoform-specific VDAC protein–lipid interface"

### Supplemental Materials and Methods

#### Recombinant protein purification

Mouse VDAC1 was a kind gift of Dr. Adam Kuszak (NIDDK, NIH). Protocols for expression, refolding, and purification of VDAC1 were based on the work of Dr. Jeff Abramson's lab (1) as described previously (2). Human VDAC1 and VDAC2 were the kind gift of Dr. Tsyr-Yan Yu (Academia Sinica, Taiwan). Human VDAC1 WT was expressed, refolded, and purified following protocol in (3). Human VDAC2 WT was expressed, refolded, and purified following the protocol in (4, 5). Human VDAC2-ΔCys was a generous gift from Dr. Radhakrishnan Mahalakshmi (Indian Institute of Science Education and Research, India) and purified as described previously (6). Human VDAC3 and VDAC3-ΔCys were the kind gifts of Dr. Lucie Bergdoll (Dr. Jeff Abramson lab, UCLA). They were expressed, refolded, and purified following the protocol in (7). hVDAC1-ΔCys was generated by mutagenesis from pET21-hVDAC1 (gift from Lucie Bergadoll) by Genewiz (Azenta Life Sciences). The plasmid sequence was confirmed by sequencing (Genewiz, Azenta Life Sciences). hVDAC1-ΔCys was expressed, refolded, and purified following the protocol in (8). Recombinant wild-type (WT) α-synuclein (αSyn) was the generous gift of Dr. Jennifer Lee (NHBLI, NIH). αSyn was expressed, purified, and characterized as described previously (9).

#### Embryonic mouse hypothalamus cell line N43/5 (mHypo43) cells

The embryonic mouse hypothalamus cell line N43/5 (mHypo43) was obtained from Cellutions Biosystems (Cedarlane, CLU127) and cultured according to the supplier's protocol. Cells were grown in DMEM (Gibco, 11965), 1% penicillin/streptomycin (Gibco, 15140), and 10% FBS (Gibco, A5256701) at 37 °C in 5% $CO_2$. Cells were passaged at 70–90% confluence using trypsin.

#### Proximity Ligation Assay (PLA)

mHypo43 were grown on 15-well µ-slides (Ibidi, 81507). PLA was carried out using Duolink In Situ reagent red (Sigma Aldrich, DUO92101) with a few modifications described here. The cells were fixed with 4% paraformaldehyde (Electron microscopy, 15710) and 0.05% glutaraldehyde (Sigma Aldrich, G5882) for 10 minutes at room temperature. The cells were then incubated with 50 mM glycine for 10 minutes. The cells were permeabilized with 0.2% Triton™ X-100 Surfact-Amps™ detergent solution (Thermofisher Scientific, 28314) in 3% BSA-PBS for 15 minutes at room temperature. After washing with PBS, the cells were incubated with Duolink blocking solution for 60 minutes at 37 °C. The cells were then incubated overnight with primary antibodies for αSyn (Cell Signaling Technology, 2628S) and COX IV (Cell Signaling Technology, 11967S) at 4 °C. Cells were washed twice with wash buffer A and incubated with anti-rabbit PLA probe plus and anti-mouse PLA probe minus along with Tom20-Alexafluor488 antibody (Proteintech, CL48866777) to label mitochondria for 1 hour at 37 °C. Cells were then incubated with Duolink ligase solution for 30 minutes, followed by Duolink amplification polymerase solution for 100 minutes at 37 °C. The cells were finally washed with wash buffer B and mounted with Duolink in situ mounting medium with DAPI (Sigma Aldrich, DUO82040). Cells were imaged with Leica TCS SP8 microscope using 100x/1.4 N.A oil objective. The cells were excited at 488 nm (Tom20-Alexa488) and 594 nm (PLA dots), and images were collected using 498–550 nm and 600–750 nm filters, respectively. The images were adjusted for contrast using ImageJ (NIH).

#### VDAC1 and VDAC2 simulation methods

Eight independent all-atom replicas of VDAC1 (2JK4 (10)) and VDAC2 (SWISS (11) model of Zebrafish VDAC2, pdb 4BUM, (12)) structures were run with the AMBER simulation package (13) for between 2 and 6 microseconds (39 aggregate microseconds for VDAC1, 36 aggregate microseconds for VDAC2), diverging according to scheduling issues on our high-performance computing resource. The forcefield was C36 (with C36m for protein parameters). The proteins were built into 25% DOPE/25% DOPC/50% DOPG bilayers consisting of 416 total lipids using the CHARMM-GUI web software (14). Simulation parameters were taken from CHARMM-GUI as appropriate for the forcefield. Temperature (310K) was maintained with a Langevin thermostat (1 ps-1 friction coefficient). PME was used for long-ranged electrostatics (12 Å cutoff with Lennard-Jones force switching between 10 Å and 12 Å). Pressure was controlled at one bar isotropic pressure using a Monte Carlo barostat. First-shell lipids were identified using the Voronoi method described in (15) and were then assigned into one of 12 angular bins around the protein, after in-plane alignment (to the reference configuration shown in Figure 9).

### Supplemental Figures

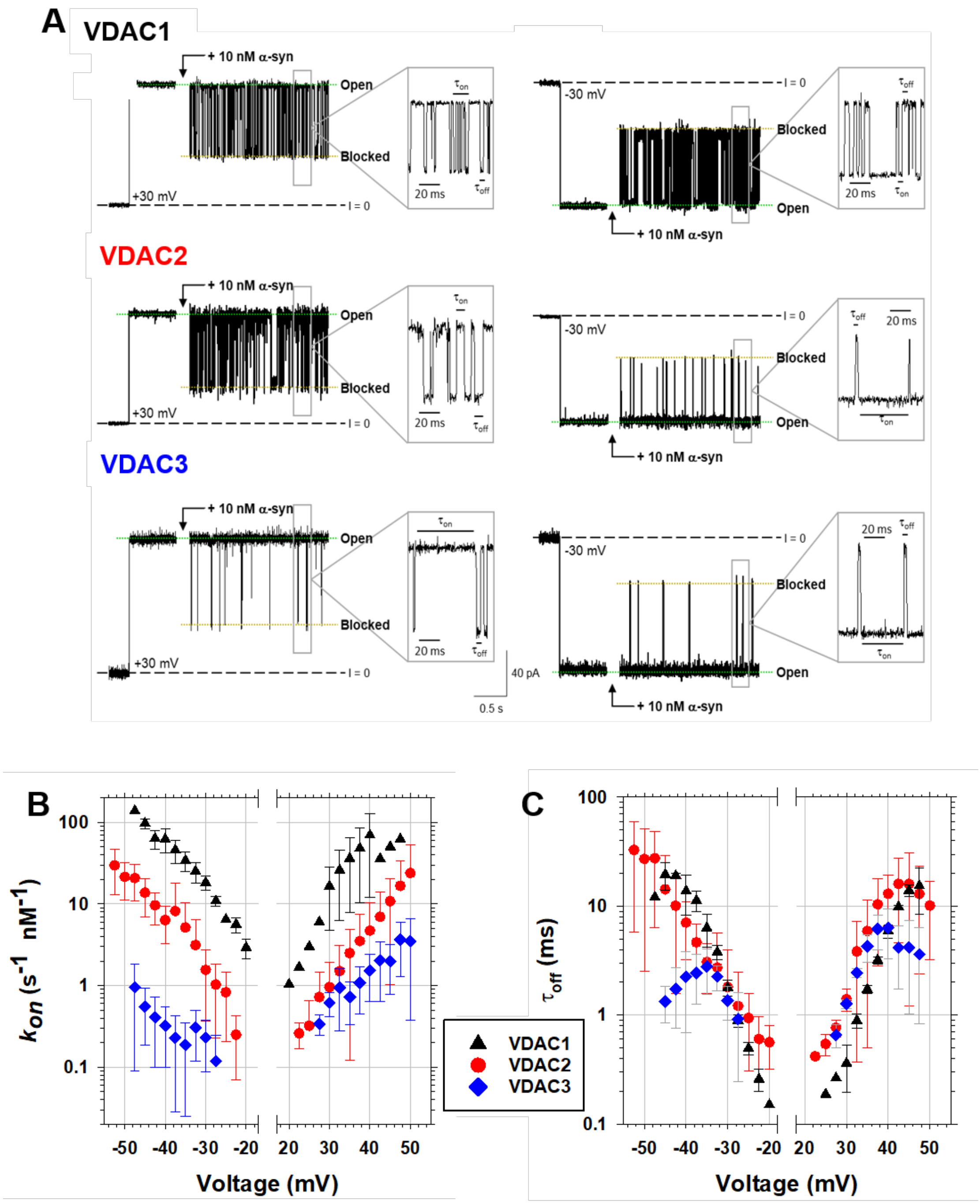


**Supplemental Figure S1.** Representative mVDAC1, hVDAC2, and hVDAC3 single-channel current traces before (*left traces*) and after addition of 10 nM αSyn (*right traces*) at ±30 mV of applied voltage. Insets show fragments of current records at a finer time scale. (C) The solid lines in (B) represent a fit to the Boltzmann equation $k_{on}$ (V) = $k_0$ exp [$n_b e|V|/(k_B T)$], where $V$ is the applied voltage, $n_b$ is the effective binding charge of 2.8 ± 0.2, and $e$, $k_B$, and $T$ have their usual meaning of elementary charge, Boltzmann constant, and absolute temperature, respectively. Data are the means of 3-7 experiments ± SD.

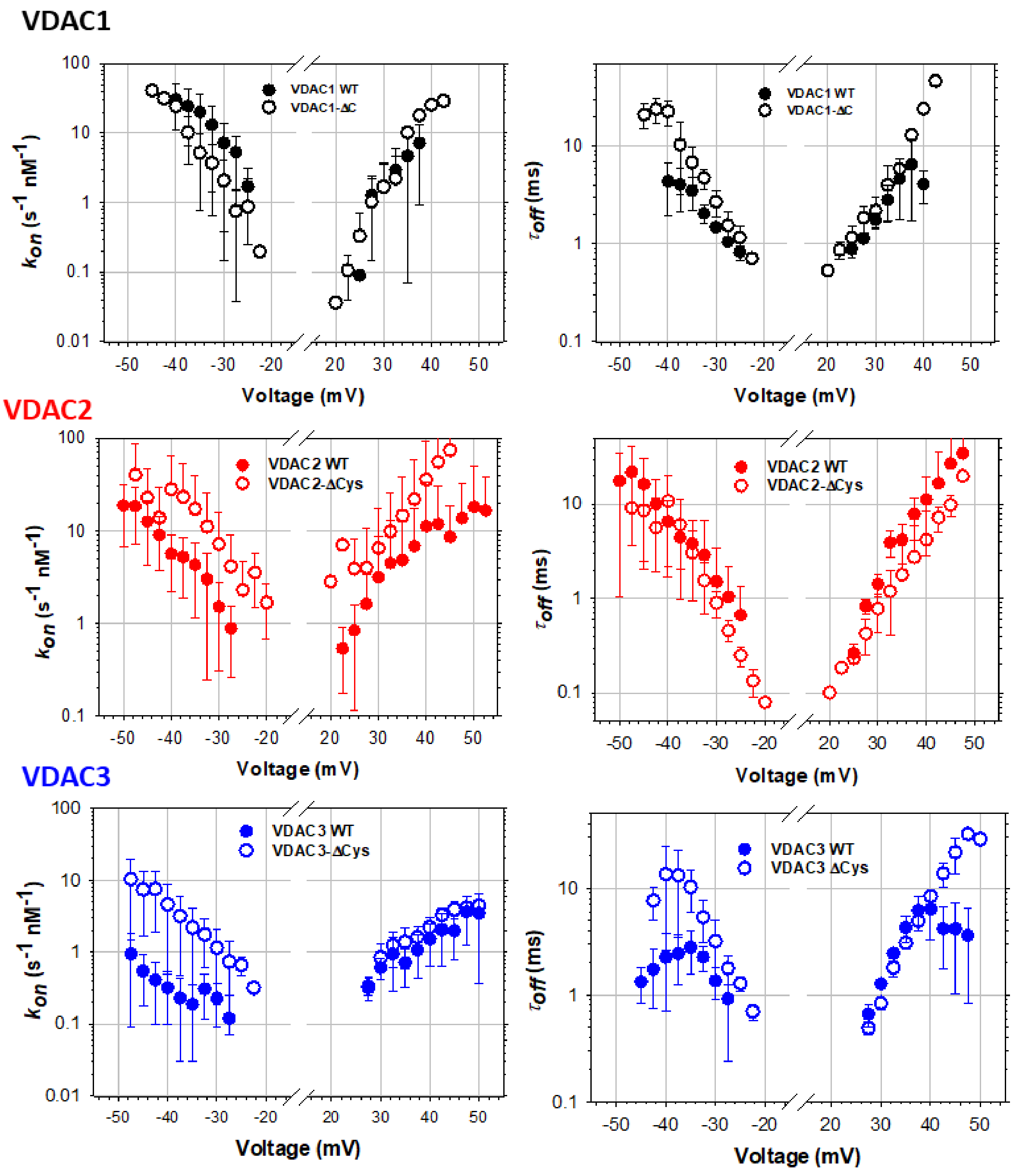


**Supplemental Figure S2. Cysteines differently affect the kinetics of αSyn interaction with VDAC isoforms.** Voltage dependences of the on-rate constants $k_{on}$ (left column) and the mean blockage time $\tau_{off}$ (right column) of αSyn binding to hVDAC1 and hVDAC1-ΔCys mutant, to hVDAC2 and hVDAC2-ΔCys mutant, and to hVDAC3 and hVDAC3-ΔCys mutant. Data are the means of 3-7 independent experiments ± SD. Experimental conditions as in Figure 5.

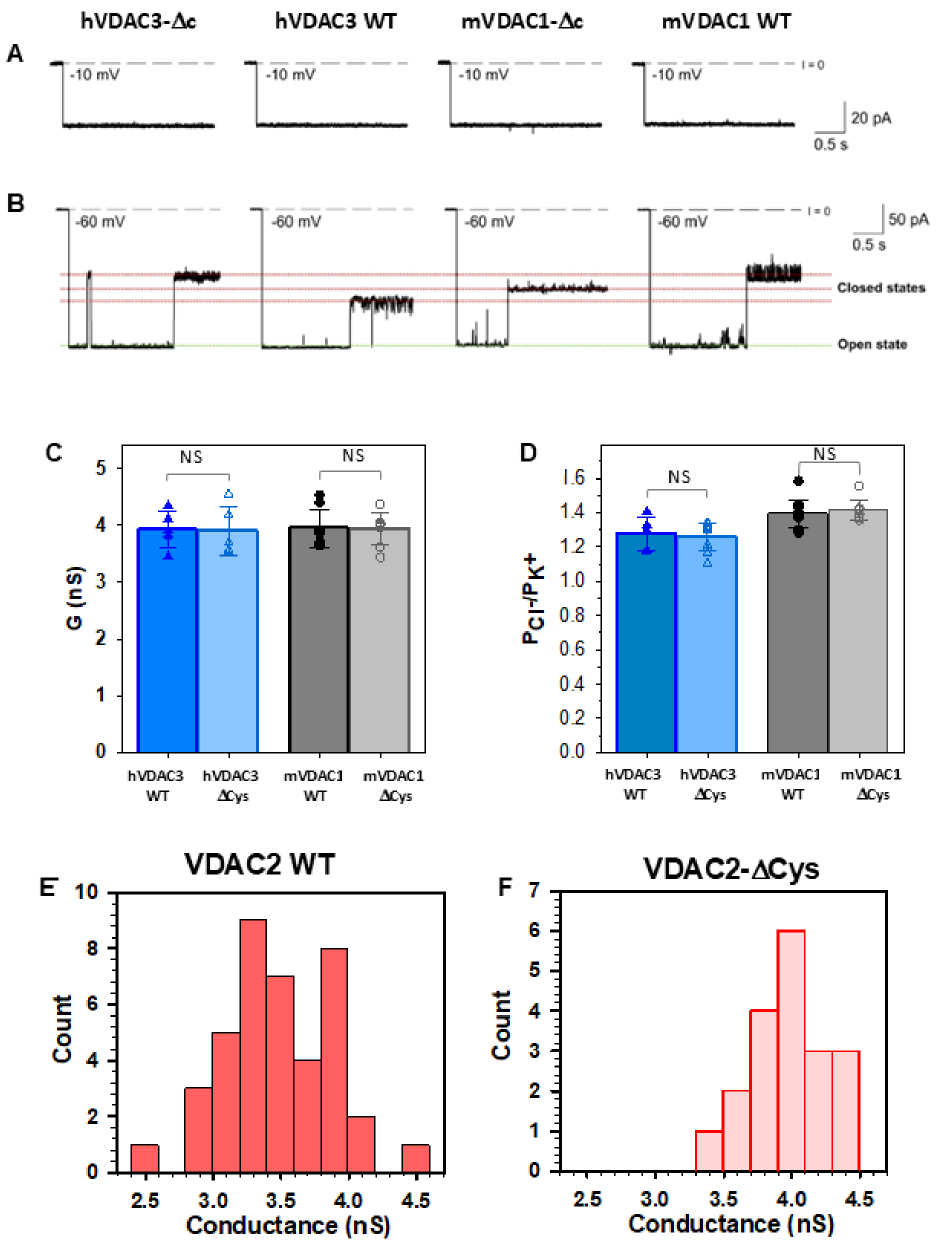


**Supplemental Figure S3. Cysteine-less hVDAC3-Δc and mVDAC1-Δc form high-conducting channels with a stable open state and voltage-induced closures typical for VDAC1 and VDAC3 WT channels.** (**A, B**) Representative single-channel current traces obtained with reconstituted hVDAC3 WT, hVDAC3-Δc, and mVDAC1 WT, mVDAC1-Δc at low (-10 mV, A) or high (-60 mV, B) applied voltages. Current records were digitally filtered at 500 Hz using a low-pass 8-pole Bessel filter. Membranes were formed from 2DOPG/DOPC/DOPE lipid mixture.

Membrane-bathing solutions consisted of 1 M KCl buffered with 5 mM HEPES at pH 7.4. (**C**, **D**) hVDAC3 and mVDAC1 cysteine-less mutants have open channel conductance and ion selectivity similar to WT. Bar graphs showing the open channel conductance (C) and ion selectivity (D) of hVDAC3 and mVDAC1, WT, and ΔCys. In (C) salt solution was 1 M KCl. In (D), the selectivity is presented as the $Cl^-$ to $K^+$ permeability ratio in 1 M/0.2 M KCl gradient. Solid circles correspond to the data points of individual experiments. Data are means of at least five independent experiments ± SD. Student's t-test with equal variance was used to check significance; NS, P > 0.5. (A-D) Adapted with permission from (7). **(E, F)** VDAC2 WT and VDAC2-ΔCys single-channel conductance histograms. The main state conductance for VDAC2 WT and VDAC2-ΔCys was measured for 24 and 5 independent experiments, respectively. Adapted with permission from (5).
